# Epistatic interactions within the influenza virus polymerase complex mediate mutagen resistance and replication fidelity

**DOI:** 10.1101/131417

**Authors:** Matthew D. Pauly, Daniel M. Lyons, Adam S. Lauring

**Author notes:** Corresponding author: Adam S. Lauring 1150 W. Medical Center Dr. MSRB1 Room 5510B Ann Arbor, MI 48109-5680.

## Abstract

Lethal mutagenesis is a broad-spectrum antiviral strategy that employs mutagenic nucleoside analogs to exploit the high mutation rate and low mutational tolerance of many RNA viruses. Studies of mutagen-resistant viruses have identified determinants of replicative fidelity and the importance of mutation rate to viral population dynamics. We have previously demonstrated the effective lethal mutagenesis of influenza virus using three nucleoside analogs as well as the virus’s high genetic barrier to mutagen resistance. Here, we investigate the mutagen-resistant phenotypes of mutations that were enriched in drug-treated populations. We find that PB1 T123A has higher replicative fitness than the wild type, PR8, and maintains its level of genome production during 5-fluorouracil treatment. Surprisingly, this mutagen-resistant variant also has an increased baseline rate of C to U and G to A mutations. A second drug-selected mutation, PA T97I, interacts epistatically with PB T123A to mediate high-level mutagen resistance, predominantly by limiting the inhibitory effect of nucleosides on polymerase activity. Consistent with the importance of epistatic interactions in the influenza polymerase, we find that nucleoside analog resistance and replication fidelity are strain dependent. Two previously identified ribavirin-resistance mutations, PB1 V43I and PB1 D27N, do not confer drug resistance in the PR8 background, and the PR8-PB1 V43I polymerase exhibits a normal baseline mutation rate. Our results highlight the genetic complexity of the influenza virus polymerase and demonstrate that increased replicative capacity is a mechanism by which an RNA virus can counter the negative effects of elevated mutation rates.

**Importance:** RNA viruses exist as genetically diverse populations. This standing genetic diversity gives them the potential to adapt rapidly, evolve resistance to antiviral therapeutics, and evade immune responses. Viral mutants with altered mutation rates or mutational tolerance have provided insights into how genetic diversity arises and how it affects the behavior of RNA viruses. To this end, we identified variants within the polymerase complex of influenza virus that are able tolerate drug-mediated increases in viral mutation rates. We find that drug resistance is highly dependent on interactions among mutations in the polymerase complex. In contrast to other viruses, influenza virus counters the effect of higher mutation rates primarily by maintaining high levels of genome replication. These findings suggest the importance of maintaining large population sizes for viruses with high mutation rates and show that multiple proteins can affect both mutation rate and genome synthesis.

## Introduction

Influenza virus remains a persistent health threat due to its rapid rate of evolution (1). This rapid rate of evolution is attributable, in part, to the virus’ very high mutation rate (2-5). Influenza virus rapidly acquires antigenic changes and antiviral resistance, which limit the effectiveness of vaccines and antiviral drugs (6, 7). As in many RNA viruses, influenza virus’ low fidelity is due to the absence of proofreading and repair mechanisms during genome replication (8-10). We have previously estimated influenza’s mutation rate to be greater than 1 × 10^−4^ mutations per nucleotide per RNA strand replicated, which suggests that approximately 2 new mutations are introduced into every newly synthesized genome (11). As a result, influenza viruses exist as swarms of distinct genetic variants, which provide a rich substrate for natural selection of adaptive mutations.

While some mutations are beneficial to a virus, the vast majority of mutations are detrimental (12). In influenza virus, we have found that 30% of single nucleotide changes are lethal and 70% decrease replicative fitness (13). Lethal mutagenesis is an antiviral strategy that utilizes nucleoside analogues to increase a virus’ mutation rate and the frequency of deleterious or lethal mutations (14, 15). The effectiveness of lethal mutagenesis has been demonstrated in many viral systems, including poliovirus, human immunodeficiency virus, foot-and-mouth disease virus (FMDV), lymphocytic choriomeningitis virus, and influenza virus (16-22). A hallmark of lethal mutagenesis is a reduction in viral specific infectivity due to the increased genesis of genomes that do not encode a functional complement of viral proteins. In addition to their mutagenic effects, many nucleoside analogs also inhibit the activity of viral RNA-dependent RNA-polymerases (RdRp) (23-26).

While initially thought to be a resistance-proof strategy, RNA virus mutants that are resistant to lethal mutagenesis have been identified by serial passage of viral populations in low concentrations of drug (20, 27-32). In most cases, mutagen resistant variants encode polymerases that exhibit increased replication fidelity, and characterization of these mutants has elucidated the molecular mechanisms governing mutation rate. With a lower baseline mutation rate, these variants require higher concentrations of mutagenic drug to increase the viral mutation rate to the same level as a wild type virus.

Other mechanisms of nucleoside resistance have been reported for RNA viruses. Polymerase mutants in FMDV mediate resistance by selecting against nucleoside misincorporation, possibly biasing the virus’ mutation spectrum (33, 34). Interestingly, a mutation in the polymerase associated 2C protein of FMDV also appears to alter mutagen-induced mutational bias (35). The DNA bacteriophage ΦX174 mitigates the impact of mutagenesis by delaying lysis and increasing its burst size, a mechanism of genetic robustness that paradoxically maintains unmutagenized progeny (36, 37). Finally, a virus’ primary sequence also affects mutagen sensitivity through its genetic robustness, or ability to buffer the fitness effects of mutations (38, 39). Collectively, these works have elucidated how mutation rate and mutational tolerance shape the diversity and structure of RNA virus populations.

Current models of lethal mutagenesis and mutagen resistance are derived almost entirely from studies of positive sense RNA viruses. In contrast to this large group of viruses, influenza virus replicates its genome using a heterotrimeric replicase complex. This complex consists of the PB2, PA, and PB1 proteins, which have 5’-cap binding, 5’-cap stealing, and RdRp activities, respectively (40). The constraints of this polymerase complex may alter the genetic barrier to mutagen resistance or its potential mechanisms. Two polymerase variants with reduced sensitivity to the mutagen ribavirin have been reported for influenza virus. The PB1 V43I mutation was identified as a minority variant upon serial passage of A/Wuhan/35/95 in low concentrations of ribavirin and appears to have altered fidelity in that genetic background (20). A second mutation, PB1 D27N, was recovered in a screen for PB1 mutants in WSN33 that maintained RNA synthesis during ribavirin treatment (41, 42).

We previously demonstrated the mutagenic activity of ribavirin, 5-azacytidine, and 5-fluorouracil in influenza virus (21). Interestingly, serial passage of influenza virus populations in sublethal concentrations of each of these three drugs did not lead to population-wide resistance. We did, however, identify three mutations – PB1 T123A, PB1 M645I, and PA T97I – that were enriched in replicate drug-selected populations (21). Here, we characterize the effects of these three mutations on nucleoside analog sensitivity, viral fitness, and replicative fidelity. Additionally, we characterize the previously identified PB1 V43I and PB1 D27N mutants in the PR8 genetic background. We show that mutagen resistant variants in the influenza polymerase complex have similar fidelity to wild type viruses. In contrast to what has been found in other viral systems, these variants mediate resistance by limiting the mutagenic and polymerase-inhibitory effects of nucleoside drugs.

## Results

We have previously shown that serial passage of influenza A PR8 in low concentrations of ribavirin, 5-fluorouracil, or 5-azacytidine does not select for population-wide resistance over 16 passages (21). This result did not preclude the possibility that there were resistance mutations present at a low level within these populations or that their phenotypic effect was masked by the impact of mutations elsewhere in the genome. Consistent with this hypothesis, next generation sequencing revealed a number of mutations that were shared among drug-passaged populations and that did not achieve fixation (21). We identified candidate resistance mutations in the PB2, PB1, and PA open reading frames based on their presence in mutagen-passaged viral populations and absence in either the starting population or control populations passaged without drug treatment. Three nonsynonymous mutations met these criteria. The PB1 T123A mutation was found in all three 5-fluorouracil (5FU) populations at frequencies of 34%, 31%, and 8%. The PA T97I mutation was found in all three ribavirin-passaged populations at frequencies of 88%, 55%, and 11%. We also identified the PB1 M645I mutation at frequencies of 90%, 14%, and 1% in these same ribavirin-passaged populations. The ribavirin resistant mutant, PB1 V43I, was not found in any of the populations, and PB1 D27N was only found in one ribavirin-passaged population at a frequency of 3%. We introduced each of these five mutations into a clean PR8 genetic background. We also generated the double mutants PB1 T123A; PA T97I and PB1 M645I; PA T97I. The PB1 M645I; PA T97I double mutant was identified in our ribavirin passaged viral populations. The PB1 T123A; PA T97I double mutant was not found naturally, but these mutations on distinct segments could plausibly interact genetically.

### Mutagen sensitivity

Given their enrichment in drug-passaged populations, we hypothesized that one or more of the mutations in the polymerase complex would mediate mutagen resistance. We tested each of the variant polymerases for reduced nucleoside sensitivity by comparing titers after replication of the corresponding virus populations in mock- or drug-treated cell cultures 24 hours post infection (Figure 1). We selected drug concentrations that would decrease infectious viral titers by 3-4 orders of magnitude so that we could observe a range of resistance phenotypes, while limiting cytotoxicity. The concentrations used – 20μM ribavirin, 20μM 5-azacytidine, and 100μM 5FU – were roughly 3 times higher than those in which our mutants were selected (21).

**Figure 1.**
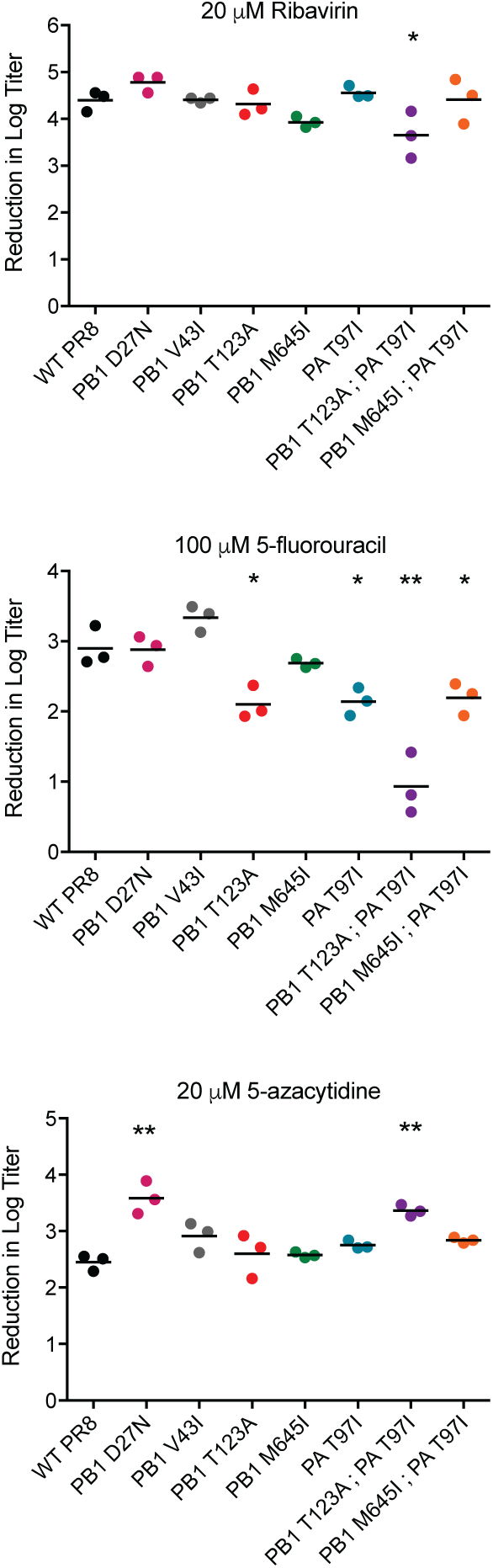
Sensitivity of influenza polymerase mutants to nucleoside analogs. MDCK cells were pretreated with media containing 0μM drug, 100μM 5-fluorouracil, 20μM ribavirin, or 20μM 5-azacytidine for 3 hours and then infected with virus at an MOI of 0.1. After 24 hours, cell free supernatants were harvested and titered by TCID_50_. The decrease in the log of the infectious titer for drug-treated samples relative to untreated samples for each virus is shown. Three replicate samples were harvested for each virus at each drug concentration and are shown along with the mean. The statistical significance of the decrease in the base 10 logarithmic titer of each mutant relative to WT PR8 was determined by one-way ANOVA with a Dunnett’s multiple comparison test. * = p-value < 0.05, ** = p-value < 0.001.

The PB1 T123A mutation, which was identified in 5FU passaged populations, conferred a 6-fold reduction in sensitivity to 100μM 5FU (Dunnett’s adjusted p-value = 0.007), but no change in sensitivity to 20μM ribavirin or 20μM 5-azacytidine. Interestingly, PA T97I, which was identified only in populations passaged in ribavirin, conferred resistance to 100μM 5FU (6-fold sensitivity reduction, Dunnett’s adjusted p-value = 0.011), but not to 20μM ribavirin or 20μM 5-azacytidine. The other mutation enriched in ribavirin-passaged populations, PB1 M645I, did not alter sensitivity to any of the three nucleoside analogs. We found that the two previously identified ribavirin resistant mutants, PB1 D27N and PB1 V43I, were just as sensitive as the wild type PR8 strain to both ribavirin and 5FU. The PB1 D27N mutant appeared to be more sensitive than WT PR8 to 5-azacytidine (13-fold increase, Dunnett’s adjusted p-value < 0.0001). These data suggest that the resistance phenotypes of PB1 D27N and PB1 V43I are dependent on strain background. Of the five single mutants tested, only PB1 T123A and PA T97I exhibited reduced sensitivity to our panel of nucleoside analogs.

The PB1 M645I and PA T97I double mutant, which was identified in ribavirin-passaged populations, exhibited a pattern of nucleoside analog sensitivity that was very similar to the PA T97I single mutant (5-fold reduction in sensitivity to 5FU, Dunnett’s adjusted p-value = 0.018). Surprisingly, the PB1 T123A; PA T97I double mutant, which was not found in passaged populations, exhibited a 93-fold reduction in sensitivity to 100μM 5FU (Dunnett’s adjusted p-value < 0.0001) and a slight, but statistically significant reduction in sensitivity to 20μM ribavirin (6-fold reduction, Dunnett’s adjusted p-value = 0.034). Paradoxically, this double mutant was more sensitive to treatment with 20μM 5-azacytidine than WT PR8 (8-fold increase, Dunnett’s adjusted p-value = 0.0004). Thus, these two mutations were synergistic with respect to 5FU resistance while at the same time increasing 5-azacytidine sensitivity.

### Replicative fitness

Mutagen resistant variants often have a fitness defect compared to their wild type counterparts. We used a serial passage competition assay to measure the replicative fitness of each mutant virus relative to the WT PR8 and performed growth curves to quantify RNA genome production. In the absence of mutagenic drug, we identified a range of fitness effects among the polymerase mutants (Figure 2A). The PB1 T123A virus was more fit than WT, even out of drug, and produced more genomes early in replication than either WT or the other mutants (Figure 2B). Both PB1 M645I and PA T97I were essentially neutral. The PB1 T123A; PA T97I double mutant, which was highly resistant to 5FU, exhibited reciprocal sign epistasis, as it had a marked decrease in fitness while each single mutant was beneficial or neutral. Consistent with the lower fitness of this mutant, it produced fewer genomes than the wild type (Figure 2B). The PB1 M645I; PA T97I double mutant had higher fitness than WT, suggesting that selection of PB1 M645I in ribavirin-passaged populations reflected culture-adaptation rather than mutagen resistance. Both PB1 D27N and PB1 V43I had dramatically reduced fitness compared with WT. The decreased fitness of PB1 V43I is consistent with previous data on its growth kinetics (20). As PB1 M645I, PB1 D27N, and the PB1 M645I; PA T97I double mutant were not resistant to any of the nucleoside drugs, we did not analyze them further.

**Figure 2.**
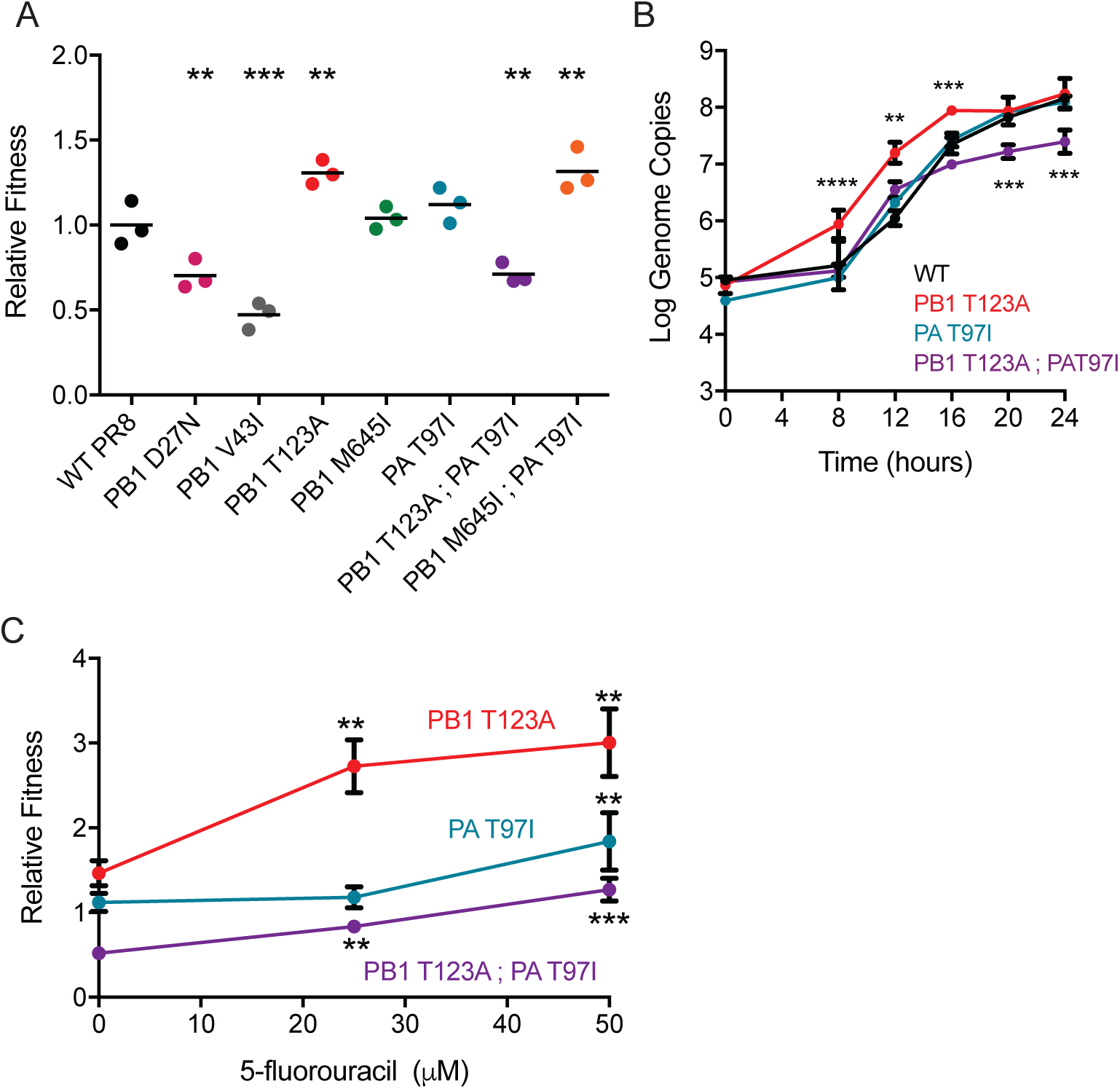
Replicative fitness of influenza polymerase mutants. (A) Direct competition assays were performed for each mutant against a WT PR8 virus containing a neutral genetic barcode. For each competition, a 1:1 starting mixture of each virus (by infectious titer) was passaged four times on MDCK cells at an MOI of 0.01. The relative changes in the amounts of the two competing viruses were determined by quantitative RT-PCR. Relative fitness was calculated as described in Methods. Data are shown for three individual competitions along with the mean. The statistical significance of the fitness values for each mutant relative to WT PR8 was determined by one-way ANOVA with a Dunnett’s multiple comparison test. * = p-value < 0.05, ** = p-value < 0.01, *** = p-value < 0.001. (B) MDCK cells were infected at an MOI of 0.1 and quantitative RT-PCR was used to measure the genome copy number. The data were analyzed using a two-way ANOVA and mutant viruses were compared with the WT using Sidak’s multiple comparison test. (C) Competition assays were performed as above for the PB1 T123A, PA T97I, and the PB1 T123A, PA T97I mutant in the presence of 5-fluorouracil. For each virus, a one-way ANOVA with a Dunnett’s multiple comparison test was used to compare the relative fitness of drug treated virus to non-treated virus. * = p-value < 0.05, ** = p-value < 0.01, *** = p-value < 0.001

We measured the fitness of PB1 T123A, PA T97I, and the double mutant PB1 T123A; PA T97I in the presence of 25μM and 50μM 5FU to better quantify their mutagen sensitivity. We used lower concentrations of drug in this serial passage experiment to avoid population extinction (see 100μM Figure 1 and (21)). We found that the relative fitness of all three 5FU resistant variants increased significantly when competed in drug, but that the double mutant only competed effectively with WT at the highest drug concentrations (Figure 2C). These data indicate that the three mutant viruses have varying resistance to 5-fluorouracil. The range of fitness values that we observed suggests that each has a different mechanism of resistance.

### Mutation rate

The most commonly identified mechanism of mutagen resistance in RNA viruses is altered baseline polymerase fidelity. We determined the baseline mutation rate of our three 5FU resistant variants using a Luria-Delbrück fluctuation test that can measure the rates of all 12 mutational classes ((11) and Figure 3). We also evaluated PB1 V43I, which has been reported to be a fidelity variant (20, 43). We used this novel fluctuation test to interrogate four of the most common mutational classes (A to G, C to U, G to A, and U to C), for which the assay has the greatest discriminatory power. These same mutational classes are also those most affected by 5FU treatment. Both PA T97I and the PB1 T123A; PA T97I double mutant had mutation rates that were very similar to those of WT. Unexpectedly, PB1 T123A made significantly more C to U and G to A mutations than WT. While we cannot rule out differences in the rates of transversion mutations, these less common mutation classes would be unlikely to affect overall polymerase fidelity. The PB1 V43I mutant had no evident resistance to nucleoside analogs or altered mutation rate in the PR8 genetic background. As the fidelity and resistance phenotypes of this mutation were demonstrated in the A/Wuhan/359/95 H3N2 strain, we suggest that its effects are modulated by epistatic interactions within the PR8 polymerase complex.

**Figure 3.**
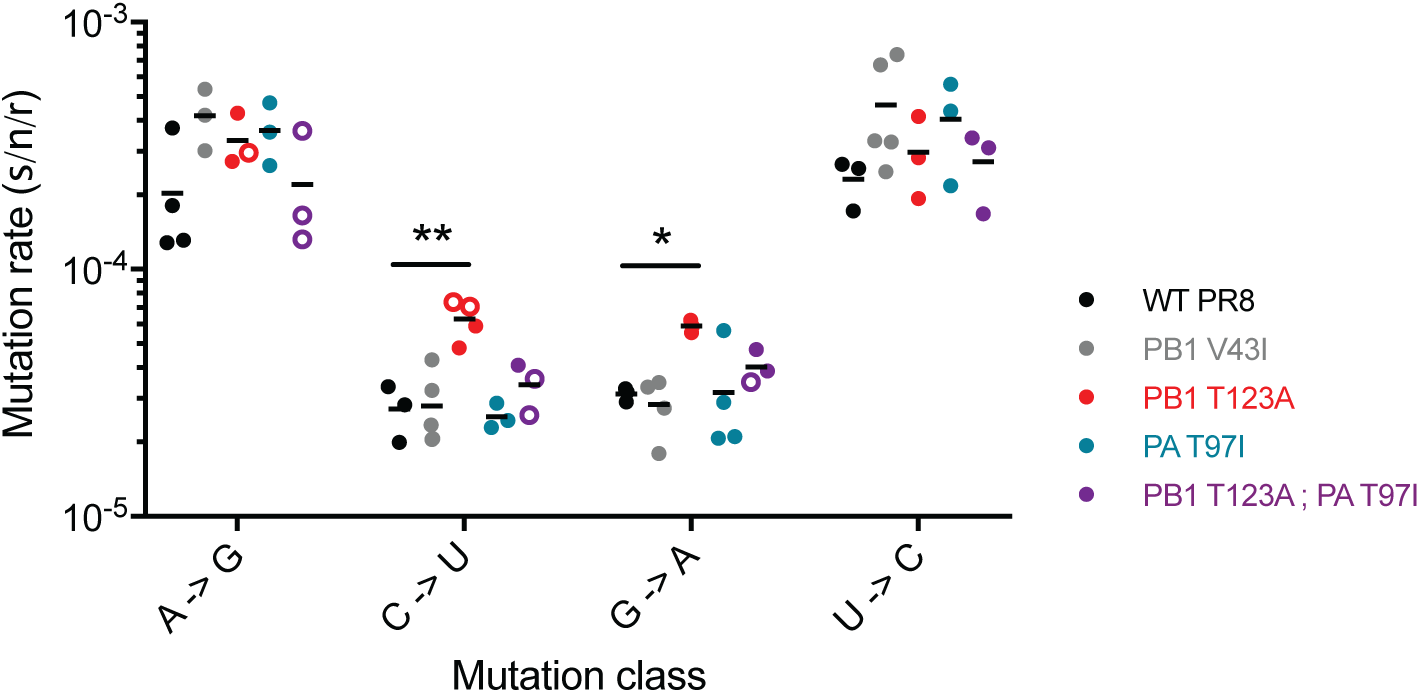
Mutation rates of influenza polymerase mutants. Rates of the four transition mutation classes were measured using a 12-class fluctuation test. Data points are color coded by virus. Measurements within the ideal P_0_ range of 0.1 - 0.7 for the null class method are shown as filled symbols. Those outside of that range, with P_0_ values between 0.7 – 0.9, are shown as open symbols. Arithmetic means are shown for the replicate measurements. The statistical significance of the differences in mutation rates for each mutation class of each mutant relative to WT PR8 was determined by one-way ANOVA with Dunnett’s multiple comparison test. * = p-value < 0.05, ** = p-value < 0.01.

We next assessed the impact of drug treatment on the mutation rates of these polymerase mutants. We used 15μM 5FU because the larger titer reductions with higher drug concentrations preclude precise measurements of mutation rates in our fluctuation test. Again, we measured the rates of the four transition mutation classes impacted by 5FU (Figure 4). The increase in transition mutations in the PA T97I mutant with 5FU treatment was similar to that of WT. In contrast, the PB1 T123A mutant, which has an increased baseline rate of C to U and G to A mutations, selectively buffers against further drug-induced increases in these same classes. This phenotype is most pronounced for C to U mutations. While 15μM 5FU increased all transition mutations approximately 5-fold in PR8, we observed almost no change in C to U mutations in the PB1 T123A mutant. The PB1 T123A; PA T97I double mutant also buffers against C to U mutations induced by 5FU, albeit not as dramatically as the PB1 T123A single mutant. Together, these data show that PB1 T123A has an increased rate for two transition mutation types while also limiting further induction by 5FU treatment.

**Figure 4.**
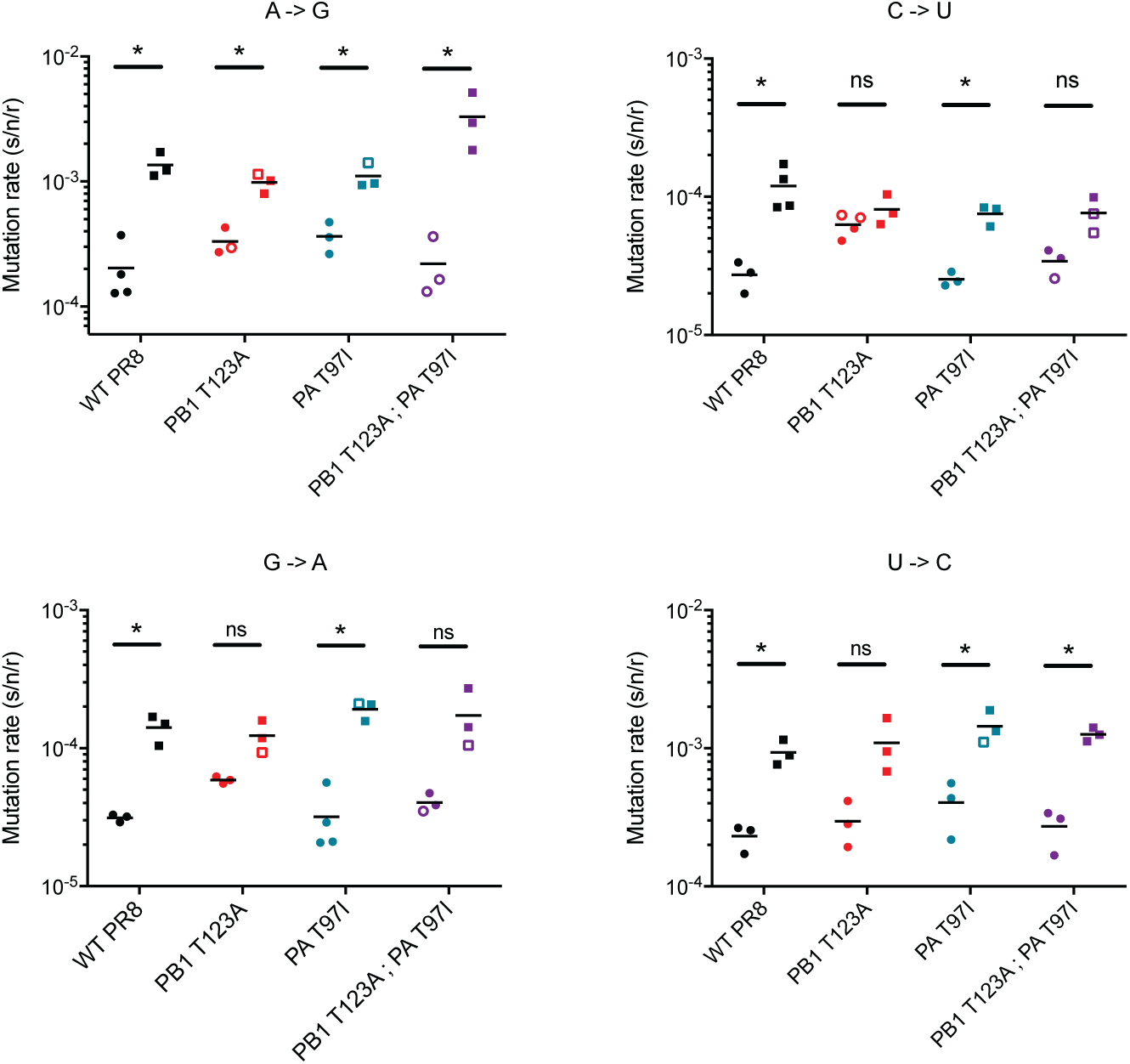
Effect of 5-fluoruracil on viral mutation rate. Mutation rates were measured as in Figure 3 using a 12-class fluctuation test. The rates of four mutation classes; A to G, C to U, G to A, and U to C, were measured in the presence of 15μM 5-fluorouracil for the indicated viruses. For each virus and mutational class, the statistical significance of the difference in mutation rates in the presence and absence of 5-fluorouracil was determined using t-tests and the Holm-Sidak correction for multiple comparisons. * = p-value < 0.05, ns = not significant.

### Genome infectivity

Increases in viral mutation rates are often accompanied by decreases in the specific infectivity of the population. Accordingly, a virus resistant to the mutagenic effects of a nucleoside analog would be expected to exhibit a smaller decrease in specific infectivity upon drug treatment. We measured the effect of 100μM 5FU on the specific infectivity of the three resistant mutants by measuring the number of infectious particles per genome (Figure 5). Drug treatment reduced the specific infectivity of all three mutants, consistent with the drug’s mutagenic effects. In all cases, the magnitude of the effect was similar to that of treated WT viruses. These data indicate that the observed decrease in 5FU-induced C to U mutations in PB1 T123A virus is not sufficient to cause a corresponding change in specific infectivity. The observed decreases in specific infectivity in 5FU are more likely due to the more common A to G and U to C mutation classes. Therefore, the limited fidelity phenotype of PB1 T123 is unlikely to contribute significantly to the virus’ mutagen resistance.

**Figure 5.**
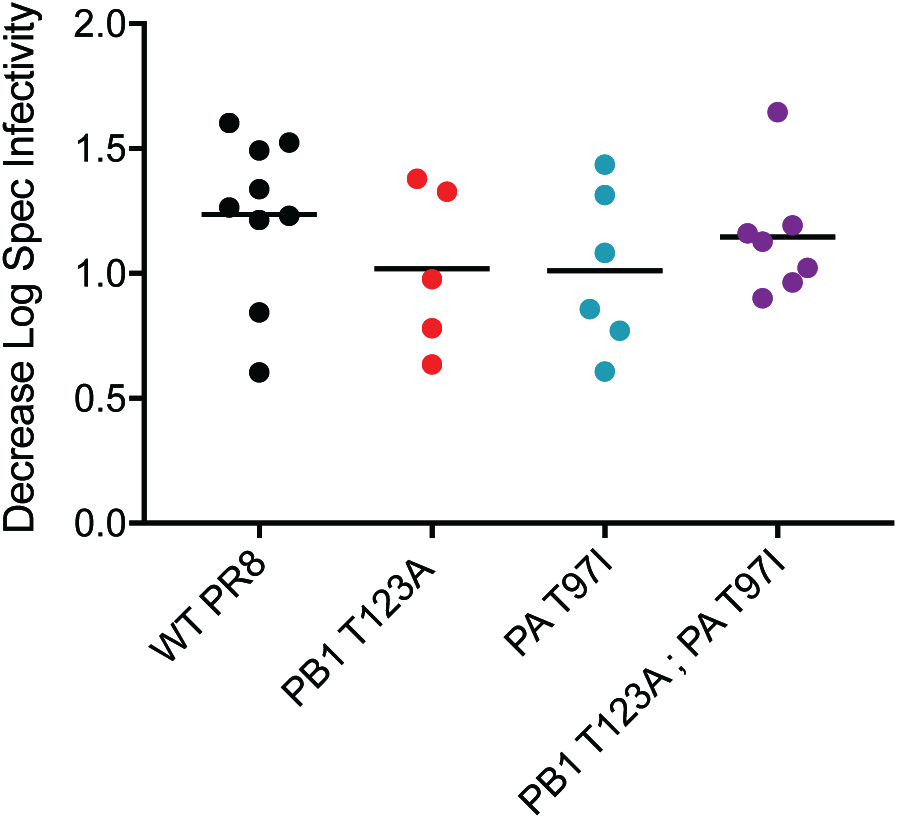
The effect of 5-fluorouracil on the specific infectivity. MDCK cells were treated with or without 100μM 5-fluorouracil and infected with influenza at an MOI of 0.1 for 24 hours. For each sample, the infectious titer was measured by TCID_50_, and the genome copy number was measured by quantitative RT-PCR. The specific infectivity was calculated as the titer divided by the genome copy number. The decreases in the log_10_ of the specific infectivity for 100μM 5-fluorouracil treated samples relative to non-treated samples are shown for replicate measurements from two or three individual experiments. There were no statistically significant differences when the data were analyzed using a one-way ANOVA.

### Genome production

Given that the resistance phenotype of our three polymerase variants was not due to altered fidelity, we evaluated their ability to resist 5FU-mediated inhibition of genome synthesis. We assessed the kinetics of genome replication in the presence and absence of 100μM 5FU by measuring the number of genomes in the supernatants of cells infected with each viral mutant. Treatment with 5FU caused a 10- to 100-fold decrease in genome copies generated by WT relative to controls (Figure 6). The PA T97I mutant exhibited a similar decrease in genome output in drug. In contrast, we observed smaller decreases in genome production in drug for both PB1 T123A and PB1 T123A; PA T97I, especially at later time points. At 24 hours post infection, there was no significant difference on the number of genomes for PB1 T123A. The PB1 T123A; PA T97I double mutant maintained its generally lower level of genome production across multiple time points, consistent with epistatic interactions between these mutations and their impact on 5FU resistance.

**Figure 6.**
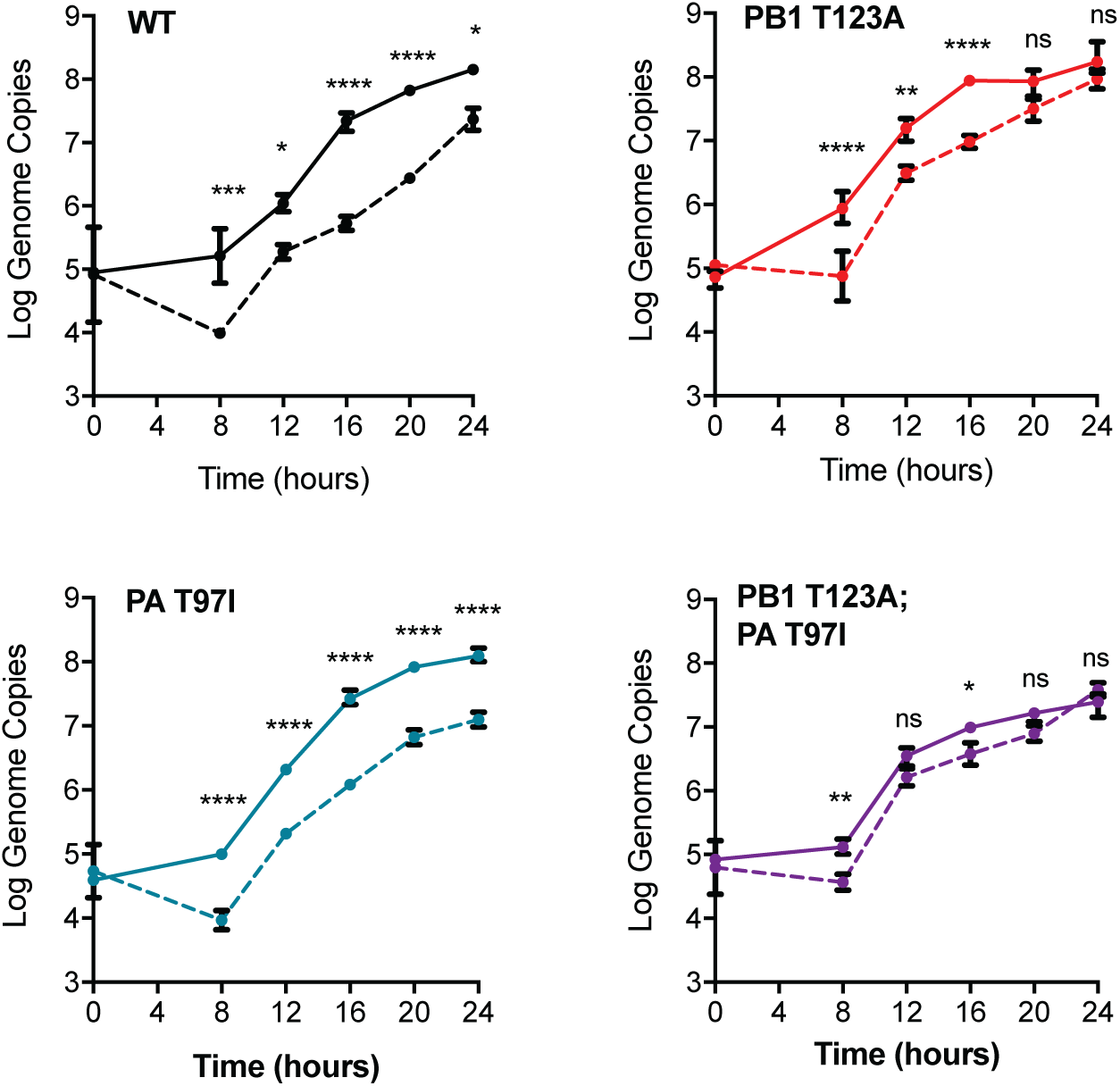
Genome production by influenza mutants during 5FU treatment. MDCK cells were infected with wild type, PB1 T123A, PA T97I, or PB1 T123A; PA T97I viruses at an MOI of 0.1 in either 0μM (solid lines) or 100μM 5-fluorouracil (dashed lines). Supernatants were collected at four hour intervals. The number of M genome segment copies per mL was determined by quantitative RT-PCR. Data are mean ± standard deviation for 3 replicates (Log_10_ scale). Genome production over time was compared in the presence and absence of drug for each virus using a two-way ANOVA with a Sidak multiple comparison test. * = p-value < 0.05, ** = p-value < 0.01, *** = p-value < 0.001, **** = p-value < 0.0001.

## Discussion

Mutagen resistant variants have been a valuable tool for probing the determinants of RNA virus mutation rates and the effect of mutation rate on viral population diversity (9, 10, 20, 27-30). We therefore investigated the mechanisms through which influenza virus can resist the antiviral effects of nucleoside analog drugs. We identified PB1 T123A and PA T97I as two 5-fluorouracil resistance mutations that interact epistatically. We also found evidence for epistasis in the previously identified ribavirin resistance mutants, PB1 D27N and PB1 V43I, as they remain sensitive to drug in the PR8 genetic background. The three mutagen-resistant viruses reported here are not high fidelity variants and the PB1 T123A variant paradoxically exhibits a higher baseline mutation rate for certain mutational classes. We identified increased genome output as the main mechanism of 5-fluroruracil resistance for PB1 T123A and resistance to drug-mediated RdRp inhibition as the mechanism for the PB1 T123A; PA T97I double mutant.

While the literature on lethal mutagenesis and fidelity variants has largely focused on viruses with a monomeric RdRp, the influenza replicase complex is composed of three proteins; PB1, PB2, and PA (44-46). The PB1 protein is the RdRp, which is shaped like a right hand with the fingers and thumb domains enclosing the palm-based active site. Mutations that alter the fidelity of RNA virus replication have rarely been observed within the active site of the RdRp (47-49). The PB1 T123 mutation is located within the fingers domain near where the RNA template enters the RdRp active site, while PA T97 is located far from the RdRp active site in the endonuclease domain of this PB1-associated protein. We suggest that the nature of the influenza polymerase complex allows for an array of intergenic, and possibly intragenic, epistatic interactions that mediate both replication fidelity and mutagen resistance.

We identified PB1 T123A as a mutation that mediates resistance to 5FU, but not ribavirin or 5-azacytidine. While mutagen selection has been used to identify high fidelity variants in a number of viral systems, the PB1 T123A virus actually has a marginally elevated mutation rate. This reduced fidelity is class-specific with the largest increase in C to U transitions. Despite its higher mutation rate, this virus had higher replicative fitness than WT and produced more genomes in the early stages of replication. We also found that while the baseline C to U mutation rate is higher for this mutant, the mutagenic effect of 5FU on this mutational class is dramatically reduced. This phenotype is similar to that of the ribavirin resistant FMDV mutant, 3D M296I (32, 50) and may reflect increased selectivity against misincorporation of 5FU. Our finding of reduced mutagenesis, however, seems inconsistent with the specific infectivity decrease we observe upon drug treatment. This may indicate that 5FU-mediated increases in the two most common mutation classes (A to G and U to C) are the main contributor to the observed decrease in specific infectivity with drug treatment. The most likely mechanism of resistance appears to be through its increase in replicative fitness in the absence of drug (Figure 2). Importantly, the mutant also maintained its genomic output in the presence of 5FU, a phenotype augmented by PA T97I (Figure 6, see below). The fact that PB1 T123A did not counter the detrimental effect of ribavirin and 5-azacytidine suggest that the resistance phenotype mediated by PB1 T123A is not broadly applicable to other nucleoside analogs.

The other mutation we investigated was located in the PB1 associated protein, PA. This mutant, PA T97I, exhibited fitness and genome production phenotypes that were very similar to those of the wild type. Interestingly, this mutant was selected during ribavirin treatment but was resistant to 5FU and not ribavirin. This mutant closely mirrored PB1 T123A in terms of its smaller decreases in infectious titer upon treatment with 5FU. Unlike PB1 T123A, the baseline and 5FU-induced mutation rates of PA T97I are similar to those of the wild type virus for all transition mutation classes. This mutant also exhibited decreases in genome production during 5FU treatment that were similar to those of WT. It is currently unclear how PA T97I mediates its resistance to 5FU. Since its resistance was more pronounced at higher concentrations of 5FU, it is possible that the technical limitations in the amount of 5FU we could use for accurate mutation rate measurements prevented us from observing a subtle phenotype.

Even though PB1 T123A and PA T97I evolved in different passage cultures, we combined them to make a double mutant. Serendipitously, we found that this double mutant exhibited the most dramatic 5FU resistance phenotype of any of the mutants we tested. These two mutations led to a reduced fitness phenotype characteristic of reciprocal sign epistasis; the combination of a mutant with increased fitness (PB1 T123A) with a neutral mutant (PA T97I) led to a double mutant with very low fitness and significantly reduced genomic RNA output. At the concentrations of 5FU used for selection of resistant variants, this double mutant has a fitness lower than wild type, even though it is highly resistant to the drug. This finding likely explains why it did not arise within our experimentally evolved populations. The double mutant virus had a nearly identical spectrum of transition mutation rates to the wild type virus and only slightly mitigated the mutagenic effect of 5FU on C to U mutations. Drug resistance seems to be driven primarily by maintaining high genomic output during 5FU treatment, an effect that appears to be more pronounced compared to the PB1 T123A single mutant (Figure 6). Treatment with nucleoside analog led to almost no reduction in the number of genome segments that are released from infected cells. This mechanism allows for more infectious viral particles to be produced than wild type despite similar levels of mutagenesis and specific infectivity decreases.

Our examination of two other mutagen-resistant variants further suggests the importance of epistatic interactions within the influenza polymerase complex. Previously, PB1 D27N was identified as a mutation that limited ribavirin inhibition of RNA synthesis in a replicon system (41, 42). We find that this mutant is not resistant to ribavirin or other mutagenic nucleoside analogs in a replication competent PR8 virus. Additionally, we found that the mutagen resistance and fidelity phenotype of PB1 V43I is strain dependent. This mutation, which mediates ribavirin resistance in the A/Wuhan/359/95 H3N2 and A/Vietnam/1203/04 H5N1 strains, is sensitive to drug in the PR8 genetic background (20). The PB1 V43I mutant, which has been suggested to be a fidelity variant even in the PR8 background (43), shows no difference in the rate of transition mutations in PR8. These findings suggest that there are likely to be epistatic interactions governing polymerase activity and fidelity in influenza virus, and that the ability of one virus to evolve resistance to a mutagen may not be reflective of how another strain evolves in the face of the same selective pressure.

While the field has often focused on the mutagenic effects of nucleoside drugs, our results suggest that their effect on viral replicative capacity may be more important in influenza virus. Both PB1 T123A and PB1 T123A; PA T97I are able to effectively maintain their titers in drug by limiting the impact of 5FU on genome production. We have previously shown that the decreases in specific infectivity associated with nucleoside analog treatment (up to 10-fold) are much less than the effect of these compounds on infectious titer output (> 1,000-fold) (21). The identification of resistant variants that maintain genome output with little impact on specific infectivity suggest that inhibition of RdRp activity is the main mechanism of action for 5FU. As we did not identify mutations that mediate ribavirin or 5-azacytidine resistance, we cannot say whether mutagenic or non-mutagenic mechanisms are dominant for these drugs.

Finally, we show that polymerases with increased replicative capacity can counteract the mutagenic effects of nucleoside drugs. This is a less recognized mechanism of mutational robustness, but one that is entirely consistent with population genetic theory (37, 51). The efficiency of negative selection is the product of the effective population size and the average mutational fitness effect. Increased genome production will lead to larger populations, and strong selection will quickly purge the large numbers of mutants with lower fitness, leaving the most fit sequence to dominate the mutant spectrum. This “safety in numbers” phenomenon leads to population-level mutational robustness even in the setting of individual-level hypersensitivity. Therefore, studies of mutagen resistance continue to provide new insights into the biochemistry of RNA virus replication and fundamental aspects of their population genetics.

## Materials and Methods

### Cells, viruses, plasmids, and drugs

Human embryonic kidney 293T fibroblasts were provided by Dr. Raul Andino (University of California San Francisco). Madin Darby canine kidney (MDCK) cells were provided by Dr. Arnold Monto (University of Michigan). Both cell lines were maintained in growth medium (Dulbecco’s modified Eagle medium (Gibco 11965) supplemented with 10% fetal bovine serum and 25 mM HEPES) at 37°C and 5% CO_2_ in a humidified incubator.

MDCK cells expressing the hemagglutinin (HA) protein of influenza A/Puerto Rico/8/1934 H1N1 (MDCK-HA cells) were generated by co-transfection with a pCABSD plasmid that expresses a Blasticidin S resistance gene and a pCAGGS plasmid encoding the influenza A/Puerto Rico/8/1934 H1N1 HA gene (52, 53). Cells stably expressing HA were selected in growth medium containing 5μg/mL Blasticidin S and were enriched for high HA expression by staining with an anti-HA antibody (1:1000 dilution, Takara c179) and an Alexa 488-conjugated anti-mouse IgG (1:200 dilution, Life Technologies A11001) followed by fluorescence-activated cell sorting on a FACSAria II (BD Biosciences). Over the course of 5 passages, cells were sorted three times to achieve a final population in which >99% of cells were positive for high level HA expression.

All eight genomic segments of influenza A/Puerto Rico/8/1934 H1N1 (PR8) (ATCC VR-1469) were cloned into the pHW2000 vector (54). Briefly, genomic RNA was harvested from the supernatants of infected cells using TRIzol reagent (Life Technologies 15596). Complementary DNA was synthesized by reverse transcription PCR using SuperScript III (Invitrogen 18080051) and Phusion high fidelity DNA polymerase (New England Biosciences M0530) with primers described by Hoffmann and colleagues (55). PCR products and pHW2000 were digested using BsmB1 (New England Biosciences R0580), Bsa1 (New England Biosciences R0535) or Aar1 (Thermo Scientific ER1581). Digested DNA was gel purified (Thermo Scientific K0691) and PCR products were ligated into pHW2000 using T4 DNA ligase (New England Biosciences M0202).

Mutant PB1 and PA segments were generated in the pHW2000 vector backbone using overlap extension PCR (56). Two rounds of PCR were performed using Phusion high fidelity DNA polymerase with pHW2000 plasmids encoding either PB1 or PA from the PR8 virus as a template, the inner mutagenic primers (PB1 D27N, Fwd 5’-CCCTTATACTGGAAACCCTCCTTACAGC-3’, Rev 5’-GCTGTAAGGAGGGTTTCCAGTATAAGGG-3’; PB1 V43I, Fwd 5’-CACCATGGATACTATCAACAGGACAC-3’, Rev 5’-GTGTCCTGTTGATAGTATCCATGGTG-3’; PB1 T123A, Fwd 5’-GTAGACAAGCTGGCACAAGGCCGAC-3’, Rev 5’-GTCGGCCTTGTGCCAGCTTGTCTAC-3’; PB1 M645I, Fwd 5’-CAATGCAGTGATAATGCCAGCACATGG-3’, Rev 5’-CCATGTGCTGGCATTATCACTGCATTG-3’; PA T97I, Fwd 5’-CAGTATTTGCAACATTACAGGGGCTGAG-3’, Rev 5’-CTCAGCCCCTGTAATGTTGCAAATACTG-3’) and the outer primers containing AarI or BsmBI restriction sites (PB1, Fwd 5’-TATTCACCTGCCTCAGGGAGCGAAAGCAGGCA-3’, Rev 5’-ATATCACCTGCCTCGTATTAGTAGAAACAAGGCATTT-3’; PA Fwd 5’-TATTCGTCTCAGGGAGCGAAAGCAGGTAC-3’, Rev 5’-ATATCGTCTCGTATTAGTAGAAACAAGGTACTT-3’). Two first round PCR reactions using Fwd inner primers with Rev outer primers and Rev inner primers with Fwd outer primers were performed. The products of these reactions were purified using a GeneJET PCR Purification Kit (Thermo K0701), mixed, and used as templates for a second-round PCR reaction using only the outer primers. Full-length PB1 and PA genes were gel purified, digested, and cloned into pHW2000 plasmid as above. PB1 containing a neutral genetic barcode was created in the same manner using the inner mutagenic primers; 5’-GATCACAACTCATTTCCAACGGAAACGGAGGGTGAGAGACAAT-3’ and 5’-ATTGTCTCTCACCCTCCGTTTCCGTTGGAAATGAGTTGTGATC-3’.

A pPOLI vector encoding enhanced green fluorescent protein (eGFP) with influenza HA packaging sequences (ΔHA-GFP) was kindly provided by Luis Martinez-Sobrido (University of Rochester). This construct contains eGFP flanked by the 78 3’-terminal bases (33 noncoding, 45 coding) and 125 5’-terminal bases (80 coding, 45 noncoding) of the influenza A/WSN/33 H1N1 HA segment, and lacks the HA translation initiation codon (57). QuikChange II site-directed mutagenesis (Agilent Technologies 200523) was used to generate twelve mutant ΔHA-GFP constructs with primers 5’-CTCGTGACCACCCTG<mutant equence>GTGCAGTGCTTCAGC-3’ and 5’-GCTGAAGCACTGCAC<mutant sequence’>CAGGGTGGTCACGAG-3’, where mutant sequence corresponds to the sequences ACCTACGAC for A to G mutation rate assessment, ACCCACGGC for C to U mutation rate assessment, ACCTGCGGC for G to A mutation rate assessment, and ATATACGGC for U to C mutation rate assessment and mutant sequence’ is its reverse complement.

Viruses were rescued from plasmid transfections of MDCK and 293T co-cultures. pHW2000 plasmids encoding all eight influenza genome segments were mixed (500ng each) in Opti-MEM (Gibco 31985062) with 8 μL of TransIT-LT1 (Mirus 2300) and incubated at room temperature for 30 minutes. Mixtures were added to 12-well plates seeded the previous day with 2 × 10^5^ 293T cells and 1 × 10^5^ MDCK cells and containing viral medium (Dulbecco’s modified Eagle medium (Gibco 11965) supplemented with 0.187% BSA, 25 mM HEPES, and 2 μg/mL TPCK treated trypsin (Worthington Biochemical 3740)). The media was changed at 24 hours, and cell free supernatants were harvested with the addition of 0.5% glycerol at 48 hours post transfection. All rescued viruses were subsequently passaged on MDCK cells at an MOI of 0.01. Passage 1 (P1) virus was harvested at 48 hours post infection. All experiments used P1 virus stocks.

Ribavirin (1-[(2*R*,3*R*,4*S*,5*R*)-3,4-dihydroxy-5-(hydroxymethyl)oxolan-2-yl]-1*H*-1,2,4-triazole-3-carboxamide) (Sigma-Aldrich R9644) was dissolved in phosphate buffered saline (PBS) at 100mM. 5-Fluorouracil (2,4-Dihydroxy-5-fluoropyrimidine) (Sigma-Aldrich F6627) was dissolved in dimethyl sulfoxide (DMSO) at 384mM. 5-Azacytidine (4-Amino-1-(β-D-ribofuranosyl)-1,3,5-triazin-2(1*H*)-one) (Sigma-Aldrich A2385) was dissolved in DMSO to make a stock at 100mM. Aliquots of these drug stocks were stored at minus 20°C.

### Mutagen sensitivity assay

Viral medium containing ribavirin, 5-azacytidine, or 5-fluorouracil was added to 24-well plates that had been seeded with 6.5 × 10^4^ MDCK cells the previous day. After three hours of drug treatment, cells were infected with 1.5 × 10^4^ pfu of virus (MOI 0.1) in 300 μl of viral media containing drug. After one hour, the inoculum was removed and 500 μl of viral medium containing drug was added. Twenty-four hours after infection, cell free supernatants were harvested by adding 0.5% glycerol, centrifuging for 5 minutes at 3,000 × g, and freezing at −80°C. Infectious viral titers were measured by TCID_50_ assay with all wells scored for cytopathic effect (CPE) at 4 days (58).

### Competition assay

Mutant PB1 or PA viruses were mixed with PR8 viruses containing a neutral genetic barcode at equivalent TCID_50_ titers. Viral mixtures were used to infect 4 × 10^5^ MDCK cells in a 12-well plate at an MOI of 0.01. After 24 hours, supernatants were harvested and passaged 3 more times on MDCK cells at an MOI of 0.01. All competitions were performed with three biological replicates. Viral RNA was harvested from the supernatants of all passages using a Purelink Pro 96 viral DNA/RNA kit (Invitrogen 12280). Superscript III and random hexamers were used to generate cDNA. Quantitative PCR was used to determine the relative amount of total PB1 (primers 5’-CAGAAAGGGGAAGATGGACA-3’ and 5’-GTCCACTCGTGTTTGCTGAA-3’), barcoded PB1 (primers 5’-ATTTCCAACGGAAACGGAGGG-3’ and 5’-AAACCCCCTTATTTGCATCC-3’), and non-barcoded PB1 (primers 5’-ATTTCCAACGGAAACGGAGGG-3’ and 5’-AAACCCCCTTATTTGCATCC-3’) in each sample. The quantities of barcoded and non-barcoded PB1 genome segments at each passage were normalized by subtracting the Ct threshold for the total PB1 primer set (ΔCt = Ct_competitior_ – Ct_total PB1_). A relative ΔCt was obtained by comparing these values at each passage to the initial P0 viral mixture (ΔΔCt = ΔCt_P1_ – ΔCt_P0_). The relative ΔCt was converted to the fold change in genome copies (Δratio = 2^−ΔΔCt^). The slope of the differences between the log_10_ Δratios of the two viruses as a function of the passage number is equal to the log_10_ relative fitness of the non-barcoded virus ([log_10_Δratio_non-barcoded_- log_10_ Δratio_barcoded_]/passage) (13).

### Specific infectivity assay

RNA was extracted from the supernatants of virally infected cells using either TRIzol Reagent (Life Technologies 15596), or Purelink Pro 96 Viral RNA/DNA Kit (Invitrogen 12280). SuperScript III (Invitrogen 18080) was used to synthesize cDNA using random hexamers. Quantitative PCR was performed on a 7500 Fast Real Time PCR system (Applied Biosystems). Superscript III RT/Platinum Taq (Thermo 2574030) was used with the primers 5’-GACCRATCCTGTCACCTCTGAC-3’ and 5’-AGGGCATTYTGGACAAAKCGTCTA −3’, and the TaqMan probe FAM-TGCAGTCCTCGCTCACTGGGCACG-3’ with Blackhole Quencher 1 with an annealing temperature of 55°C for M segment copy number measurement. Quantification of cDNA copy number based on cycle threshold (Ct) values was performed using standard curves from ten-fold dilutions of plasmid containing the M gene of A/Puerto Rico/8/1934 H1N1. The ratio of the infectious titer per mL to the genome copy number per mL is the specific infectivity of the sample.

### Mutation rate assay

Twenty-four wells containing 1.2 × 10^4^ MDCK-HA cells were infected with 400 TCID_50_ of influenza viruses encoding mutant ΔHA-GFP segments in viral medium. Supernatants were transferred to black 96-well plates (Perkin Elmer 6005182) containing 1.5 × 10^4^ MDCK cells and 50μL of viral media at 17-23 hours post infection, depending on the mutation class and drug treatment. Two wells were infected with virus equivalent to the amount used to initially infect the parallel cultures. These wells were used to determine N_i_ (initial population size) in the mutation rate calculation. After 14 hours, cells were fixed using 2% formaldehyde for 20 minutes. Cells were rinsed with PBS and permeabilized using 0.1% triton-X-100 for 8 minutes. After rinsing again, nonspecific antibody binding sites were blocked using 2% BSA in PBS containing 0.1% tween-20 (PBS-T) for 1 hour. Cells were stained with 1:5000 Hoechst (Life Technologies 33342) and 1:400 anti-GFP Alexa 647 conjugate (Life Technologies A31852) diluted in 2% BSA in PBS-T for 1 hour. After three washes with PBS-T the plates were sealed with black tape prior to removing the final wash. Plates were imaged using an ImageXpress Micro (Molecular Dynamics) using DAPI, Cy5, and FITC specific filter cubes with a 4x magnification lens. The entire surface area of each well was imaged using four non-overlapping quadrants. MetaXpress version 6 software (Molecular Dynamics) was used to count cellular nuclei and antibody stained cells. Cells expressing fluorescent GFP were manually counted from the collected images (11).

A null class Luria-Delbrück fluctuation test was used to calculate the mutation rates with the equation μ_(s/n/r)_ = -ln(P_0_)/(N_f_-N_i_), where μ_(s/n/r)_ is the mutation rate per strand replicated, P_0_ is the proportion of cultures that do not contain a cell infected by a virus encoding fluorescent eGFP, and N_f_ and N_i_ are the final and initial viral population sizes, as determined by anti-GFP antibody staining (59, 60). If the number of green cells in a culture was greater than 0.8 (N_f_/N_i_) it was removed from the calculation because it likely contained a pre-existing fluorescent revertant in the inoculum. Cultures with this many green cells were extremely rare due to the use of a small inoculum (N_i_). The null class fluctuation test measurement is most precise when P_0_ is between 0.1 and 0.7. As a result of lower titers from drug treated viral cultures, not all of our measurements fell within this range. Replicates for which the P_0_ was greater than 0.7 are indicated in the mutation rate figures (Figures 3 and 4)

## Acknowledgments

We thank Will Fitzsimmons for technical assistance and Robert Woods for helpful suggestions. This work was supported by a Clinician Scientist Development Award from the Doris Duke Charitable Foundation (CSDA 2013105) and R01 AI118886, both to ASL. MDP was supported by the Michigan Predoctoral Training Program in Genetics (T32GM007544) and an EDGE award from the University of Michigan Endowment for Basic Sciences. DML was supported by University of Michigan Medical Scientist Training Program (T32GM007863). The funders had no role in study design, data collection and interpretation, or the decision to submit the work for publication.

## References

1. Nelson MI, Holmes EC. 2007. The evolution of epidemic influenza. Nat Rev Genet 8:196–205.

2. Parvin JD, Moscona A, Pan WT, Leider JM, Palese P. 1986. Measurement of the mutation rates of animal viruses: influenza A virus and poliovirus type 1. J Virol 59:377–383.

3. Nobusawa E, Sato K. 2006. Comparison of the mutation rates of human influenza A and B viruses. J Virol 80:3675–3678.

4. Suárez-López P, Ortín J. 1994. An estimation of the nucleotide substitution rate at defined positions in the influenza virus haemagglutinin gene. J Gen Virol 75 (Pt 2):389–393.

5. Suárez P, Valcárcel J, Ortín J. 1992. Heterogeneity of the mutation rates of influenza A viruses: isolation of mutator mutants. J Virol 66:2491–2494.

6. Paules C, Subbarao K. 2017. Influenza. Lancet.

7. Osterholm MT, Kelley NS, Sommer A, Belongia EA. 2012. Efficacy and effectiveness of influenza vaccines: a systematic review and meta-analysis. Lancet Infect Dis 12:36–44.

8. Steinhauer DA, Domingo E, Holland JJ. 1992. Lack of evidence for proofreading mechanisms associated with an RNA virus polymerase. Gene 122:281–288.

9. Smith EC, Blanc H, Vignuzzi M, Denison MR. 2013. Coronaviruses Lacking Exoribonuclease Activity Are Susceptible to Lethal Mutagenesis: Evidence for Proofreading and Potential Therapeutics. PLoS Pathog 9:e1003565.

10. Smith EC, Sexton NR, Denison MR. 2014. Thinking Outside the Triangle: Replication Fidelity of the Largest RNA Viruses. Annual Review of Virology 1:111–132.

11. Pauly MD, Procario M, Lauring AS. 2017. The mutation rates and mutational bias of influenza A virus. bioRxiv.

12. Sanjuán R. 2010. Mutational fitness effects in RNA and single-stranded DNA viruses: common patterns revealed by site-directed mutagenesis studies. Philos Trans R Soc Lond, B, Biol Sci 365:1975–1982.

13. Visher E, Whitefield SE, McCrone JT, Fitzsimmons W, Lauring AS. 2016. The Mutational Robustness of Influenza A Virus. PLoS Pathog 12:e1005856–25.

14. Anderson JP, Daifuku R, Loeb LA. 2004. Viral error catastrophe by mutagenic nucleosides. Annu Rev Microbiol 58:183–205.

15. Bull JJ, Sanjuán R, Wilke CO. 2007. Theory of lethal mutagenesis for viruses. J Virol 81:2930–2939.

16. Lee CH, Gilbertson DL, Novella IS, Huerta R, Domingo E, Holland JJ. 1997. Negative effects of chemical mutagenesis on the adaptive behavior of vesicular stomatitis virus. J Virol 71:3636–3640.

17. Crotty S, Cameron CE, Andino R. 2001. RNA virus error catastrophe: direct molecular test by using ribavirin. Proc Natl Acad Sci USA 98:6895–6900.

18. Loeb LA, Mullins JI. 2000. Lethal mutagenesis of HIV by mutagenic ribonucleoside analogs. AIDS Res Hum Retroviruses 16:1–3.

19. Grande-Pérez A, Sierra S, Castro MG, Domingo E, Lowenstein PR. 2002. Molecular indetermination in the transition to error catastrophe: systematic elimination of lymphocytic choriomeningitis virus through mutagenesis does not correlate linearly with large increases in mutant spectrum complexity. Proc Natl Acad Sci USA 99:12938–12943.

20. Cheung PPH, Watson SJ, Choy K-T, Fun Sia S, Wong DDY, Poon LLM, Kellam P, Guan Y, Malik Peiris JS, Yen H-L. 2014. Generation and characterization of influenza A viruses with altered polymerase fidelity. Nature Communications 5:4794.

21. Pauly MD, Lauring AS. 2015. Effective lethal mutagenesis of influenza virus by three nucleoside analogs. J Virol 89:3584–3597.

22. Baranovich T, Wong S-S, Armstrong J, Marjuki H, Webby RJ, Webster RG, Govorkova EA. 2013. T-705 (favipiravir) induces lethal mutagenesis in influenza A H1N1 viruses in vitro. J Virol 87:3741–3751.

23. Graham AF, Kirk C. 1965. Effect of 5-fluorouracil on the growth of bacteriophage R17. J Bacteriol 90:928–935.

24. Furuta Y, Takahashi K, Kuno-Maekawa M, Sangawa H, Uehara S, Kozaki K, Nomura N, Egawa H, Shiraki K. 2005. Mechanism of action of T-705 against influenza virus. Antimicrob Agents Chemother 49:981–986.

25. Vanderlinden E, Vrancken B, Van Houdt J, Rajwanshi VK, Gillemot S, Andrei G, Lemey P, Naesens L. 2016. Distinct Effects of T-705 (Favipiravir) and Ribavirin on Influenza Virus Replication and Viral RNA Synthesis. Antimicrob Agents Chemother 60:6679–6691.

26. Eriksson B, Helgstrand E, Johansson NG, Larsson A, Misiorny A, Norén JO, Philipson L, Stenberg K, Stening G, Stridh S, Oberg B. 1977. Inhibition of influenza virus ribonucleic acid polymerase by ribavirin triphosphate. Antimicrob Agents Chemother 11:946–951.

27. Pfeiffer JK, Kirkegaard K. 2003. A single mutation in poliovirus RNA-dependent RNA polymerase confers resistance to mutagenic nucleotide analogs via increased fidelity. Proc Natl Acad Sci USA 100:7289–7294.

28. Vignuzzi M, Stone JK, Arnold JJ, Cameron CE, Andino R. 2006. Quasispecies diversity determines pathogenesis through cooperative interactions in a viral population. Nature 439:344–348.

29. Coffey LL, Beeharry Y, Bordería AV, Blanc H, Vignuzzi M. 2011. Arbovirus high fidelity variant loses fitness in mosquitoes and mice. Proceedings of the National Academy of Sciences 108:16038–16043.

30. Zeng J, Wang H, Xie X, Yang D, Zhou G, Yu L. 2013. An increased replication fidelity mutant of foot-and-mouth disease virus retains fitness in vitro and virulence in vivo. Antiviral Res 100:1–7.

31. Zeng J, Wang H, Xie X, Li C, Zhou G, Yang D, Yu L. 2014. Ribavirin-Resistant Variants of Foot-and-Mouth Disease Virus: the Effect of Restricted Quasispecies Diversity on Viral Virulence. J Virol 88:4008–4020.

32. Sierra M, Airaksinen A, González-López C, Agudo R, Arias A, Domingo E. 2007. Foot-and-mouth disease virus mutant with decreased sensitivity to ribavirin: implications for error catastrophe. J Virol 81:2012–2024.

33. Agudo R, Ferrer-Orta C, Arias A, la Higuera de I, Perales C, Pérez-Luque R, Verdaguer N, Domingo E. 2010. A multi-step process of viral adaptation to a mutagenic nucleoside analogue by modulation of transition types leads to extinction-escape. PLoS Pathog 6:e1001072.

34. Perales C, Agudo R, Domingo E. 2009. Counteracting quasispecies adaptability: extinction of a ribavirin-resistant virus mutant by an alternative mutagenic treatment. PLoS ONE 4:e5554.

35. Agudo R, la Higuera de I, Arias A, Grande-Pérez A, Domingo E. 2016. Involvement of a joker mutation in a polymerase-independent lethal mutagenesis escape mechanism. Virology 494:257–266.

36. Pereira-Gómez M, Sanjuán R. 2014. Delayed lysis confers resistance to the nucleoside analogue 5-fluorouracil and alleviates mutation accumulation in the single-stranded DNA bacteriophage X174. J Virol 88:5042–5049.

37. Elena SF. 2012. RNA virus genetic robustness: possible causes and some consequences. Curr Opin Virol 2:525–530.

38. Graci JD, Gnädig NF, Galarraga JE, Castro C, Vignuzzi M, Cameron CE. 2012. Mutational robustness of an RNA virus influences sensitivity to lethal mutagenesis. J Virol 86:2869–2873.

39. Lauring AS, Acevedo A, Cooper SB, Andino R. 2012. Codon usage determines the mutational robustness, evolutionary capacity, and virulence of an RNA virus. Cell Host and Microbe 12:623–632.

40. Velthuis Te AJW, Fodor E. 2016. Influenza virus RNA polymerase: insights into the mechanisms of viral RNA synthesis. Nat Rev Micro 14:479–493.

41. Binh NT, Wakai C, Kawaguchi A, Nagata K. 2014. Involvement of the N-terminal portion of influenza virus RNA polymerase subunit PB1 in nucleotide recognition. Biochem Biophys Res Commun 443:975–979.

42. Binh NT, Wakai C, Kawaguchi A, Nagata K. 2013. The N-terminal region of influenza virus polymerase PB1 adjacent to the PA binding site is involved in replication but not transcription of the viral genome. Front Microbiol 4:398.

43. Naito T, Mori K, Ushirogawa H, Takizawa N, Nobusawa E, Odagiri T, Tashiro M, Ohniwa RL, Nagata K, Saito M. 2017. Generation of a Genetically Stable High-Fidelity Influenza Vaccine Strain. J Virol 91:e01073–16.

44. Velthuis Te AJW, Fodor E. 2016. Influenza virus RNA polymerase: insights into the mechanisms of viral RNA synthesis. Nat Rev Micro 14:479–493.

45. Pflug A, Guilligay D, Reich S, Cusack S. 2014. Structure of influenza A polymerase bound to the viral RNA promoter. Nature 516:355–360.

46. Reich S, Guilligay D, Pflug A, Malet H, Berger I, Crépin T, Hart D, Lunardi T, Nanao M, Ruigrok RWH, Cusack S. 2014. Structural insight into cap-snatching and RNA synthesis by influenza polymerase. Nature 516:361–366.

47. Arnold JJ, Vignuzzi M, Stone JK, Andino R, Cameron CE. 2005. Remote site control of an active site fidelity checkpoint in a viral RNA-dependent RNA polymerase. J Biol Chem 280:25706–25716.

48. Korneeva VS, Cameron CE. 2007. Structure-function relationships of the viral RNA-dependent RNA polymerase: fidelity, replication speed, and initiation mechanism determined by a residue in the ribose-binding pocket. J Biol Chem 282:16135–16145.

49. Campagnola G, McDonald S, Beaucourt S, Vignuzzi M, Peersen OB. 2014. Structure-Function Relationships Underlying the Replication Fidelity of Viral RNA-Dependent RNA Polymerases. J Virol 89:275–286.

50. Arias A, Arnold JJ, Sierra M, Smidansky ED, Domingo E, Cameron CE. 2008. Determinants of RNA-dependent RNA polymerase (in)fidelity revealed by kinetic analysis of the polymerase encoded by a foot-and-mouth disease virus mutant with reduced sensitivity to ribavirin. J Virol 82:12346–12355.

51. Lauring AS, Frydman J, Andino R. 2013. The role of mutational robustness in RNA virus evolution. Nat Rev Micro 11:327–336.

52. Marsh GA, Hatami R, Palese P. 2007. Specific residues of the influenza A virus hemagglutinin viral RNA are important for efficient packaging into budding virions. J Virol 81:9727–9736.

53. Baker SF, Nogales A, Finch C, Tuffy KM, Domm W, Perez DR, Topham DJ, Martínez-Sobrido L. 2014. Influenza A and B Virus Intertypic Reassortment through Compatible Viral Packaging Signals. J Virol 88:10778–10791.

54. Hoffmann E, Neumann G, Kawaoka Y, Hobom G, Webster RG. 2000. A DNA transfection system for generation of influenza A virus from eight plasmids. Proc Natl Acad Sci USA 97:6108–6113.

55. Hoffmann E, Stech J, Guan Y, Webster RG, Perez DR. 2001. Universal primer set for the full-length amplification of all influenza A viruses. Arch Virol 146:2275–2289.

56. Ho SN, Hunt HD, Horton RM, Pullen JK, Pease LR. 1989. Site-directed mutagenesis by overlap extension using the polymerase chain reaction. Gene 77:51–59.

57. Martínez-Sobrido L, Cadagan R, Steel J, Basler CF, Palese P, Moran TM, García-Sastre A. 2010. Hemagglutinin-pseudotyped green fluorescent protein-expressing influenza viruses for the detection of influenza virus neutralizing antibodies. J Virol 84:2157–2163.

58. Reed LJ, Muench H. 1938. A SIMPLE METHOD OF ESTIMATING FIFTY PER CENT ENDPOINTS, 27:493–497.

59. Furió V, Moya A, Sanjuán R. 2005. The cost of replication fidelity in an RNA virus. Proc Natl Acad Sci USA 102:10233–10237.

60. Foster PL. 2006. Methods for determining spontaneous mutation rates. Meth Enzymol 409:195–213.

